# Eicosatetraynoic Acid Regulates Pro-Fibrotic Pathways in an Induced Pluripotent Stem Cell Derived Macrophage:Human Intestinal Organoid Model of Crohn’s Disease

**DOI:** 10.1101/2024.01.30.577959

**Authors:** Ingrid Jurickova, Benjamin W. Dreskin, Elizabeth Angerman, Erin Bonkowski, Kentaro Tominaga, Kentaro Iwasawa, Tzipi Braun, Takanori Takebe, Michael A. Helmrath, Yael Haberman, James M. Wells, Lee A. Denson

## Abstract

**Background and Aims:** We previously identified small molecules predicted to reverse an ileal gene signature for future Crohn’s Disease (CD) strictures. Here we used a new human intestinal organoid (HIO) model system containing macrophages to test a lead candidate, eicosatetraynoic acid (ETYA).

**Methods:** Induced pluripotent stem cell lines (iPSC) were derived from CD patients and differentiated into macrophages and HIOs. Macrophages and macrophage:HIO co-cultures were exposed to lipopolysaccharide (LPS) with and without ETYA pre-treatment. Cytospin and flow cytometry characterized macrophage morphology and activation markers, and RNA sequencing defined the global pattern of macrophage gene expression. TaqMan Low Density Array, Luminex multiplex assay, immunohistologic staining, and sirius red polarized light microscopy were performed to measure macrophage cytokine production and HIO pro-fibrotic gene expression and collagen content.

**Results:** iPSC-derived macrophages exhibited morphology similar to primary macrophages and expressed inflammatory macrophage cell surface markers including CD64 and CD68. LPS-stimulated macrophages expressed a global pattern of gene expression enriched in CD ileal inflammatory macrophages and matrisome secreted products, and produced cytokines and chemokines including CCL2, IL1B, and OSM implicated in refractory disease. ETYA suppressed CD64 abundance and pro-fibrotic gene expression pathways in LPS stimulated macrophages. Co-culture of LPS-primed macrophages with HIO led to up-regulation of fibroblast activation genes including *ACTA2* and *COL1A1*, and an increase in HIO collagen content. ETYA pre-treatment prevented pro-fibrotic effects of LPS-primed macrophages.

**Conclusions:** ETYA inhibits pro-fibrotic effects of LPS-primed macrophages upon co-cultured HIO. This model may be used in future untargeted screens for small molecules to treat refractory CD.

## Introduction

A recent adult-onset ileal Crohn’s Disease (CD) single cell RNA sequencing (scRNASeq) analysis implicated an inflammatory macrophage:fibroblast circuit in anti-TNF non-response for children with CD enrolled in the RISK pediatric CD inception cohort study.^1,2^ This report linked inflammatory macrophage production of TNFA, IL1B, and OSM, and activated fibroblast production of the CCL2 and CCL7 chemokines, to anti-TNF non-response. A subsequent report utilized the Olink proximity extension assay to identify circulating protein biomarkers of a refractory course for children in RISK.^3^ This defined novel secreted factors including IL7, MMP10, IL12B, and CCL11 associated with stricturing complications, and TNFSF14, CCL4, IL15RA, TNFB, and CD40 associated with internal penetrating complications. Ileal scRNASeq confirmed that each of the factors predictive of stricturing, and IL15RA, CCL4, and CD40 associated with internal penetrating complications, were expressed by inflammatory macrophages and/or activated fibroblasts.^1,3^ These studies have suggested that inflammatory macrophage:fibroblast cross-talk is a critical mechanism of refractory CD, and have implicated specific cytokines and chemokines.

We completed a perturbagen analysis to prioritize small molecules likely to reverse the pro-inflammatory and pro-fibrotic ileal gene expression signature observed in pediatric CD patients who did not respond to anti-TNF and subsequently developed strictures.^4^ Our CMap connectivity analysis identified 140 small molecules predicted to reverse inflammatory and fibrotic pathways linked to anti-TNF response and CD strictures. The synthetic long chain fatty acid eicosatetraynoic acid (ETYA) was predicted to inhibit macrophage activation and fibroblast proliferation. Induced pluripotent stems cell (iPSC) derived human intestinal organoids (HIO) which contain both epithelial cells and fibroblasts may be used to explore genetic and environmental mechanisms of CD pathogenesis.^5-7^ We recently reported that both ETYA, and the microbial metabolite butyrate, reduced tissue collagen content in CD patient iPSC-derived HIO.^8^

A limitation of iPSC-derived HIO in modeling diseases including CD has been the lack of tissue resident immune cells.^9^ This has been addressed by the development of methods to also differentiate immune cells including macrophages from iPSC lines.^10^ However, whether macrophages differentiated from CD patient-derived iPSC lines could be used to study pathways implicated in tissue fibrosis was not known. In the current study, we developed a CD iPSC-derived macrophage:HIO co-culture system to test effects of the microbial product LPS, and the long-chain fatty acid ETYA, on both inflammatory and fibrotic pathways implicated in refractory CD. We confirmed that LPS stimulated macrophages from CD patients were enriched for an inflammatory and pro-fibrotic gene expression program similar to CD patients, and that ETYA prevented development of the pro-fibrotic phenotype observed in macrophage:HIO co-culture.^1,4^

## Materials and Methods

### Macrophage generation and characterization

Peripheral blood mononuclear cells (PBMC) were obtained from four pediatric-onset CD patients. Induced pluripotent stem cell (iPSC) lines (Lines A, B, C, and D) were derived from donor PBMC as previously described.^9,11^ The iPSC lines completed quality control testing for mycoplasma, karyotype, Short Tandem Repeat/Electropherogram, stemness analysis, and functional pluripotency. Macrophages were generated from iPSC using the STEMdiff™ Hematopoietic Kit (StemCell Technologies Cambridge, MA) and differentiation protocol as reported.^10^ At day -1 iPSC were prepared in mTeSR™1 (StemCell Technologies Cambridge, MA) and plated at approximately 40 colonies per well on Matrigel pre-coated wells. On day 12 a suspension of hematopoietic progenitor cells was harvested for downstream assays. Hematopoietic progenitor cells were re-suspended at 0.5 x10^6^ in 2 mL media: DMEM (ThermoFisher Waltham, MA), 10% FBS (Corning Corning, NY), 1% P/S (ThermoFisher Waltham, MA), 10 ng/mL rhGM-CSF (R&D Systems Minneapolis, MN) and 25 ng/mL rhM-CSF (R&D Systems Minneapolis, MN). On day 13 adherent macrophages were harvested using Trypsin-EDTA (0.25%) (ThermoFisher Waltham, MA). Macrophages were characterized by cytospin and flow cytometry under basal conditions and following LPS (Sigma-Aldrich St. Louis, MO) stimulation (100 ng/nL for 72 hours) with CD14 (R&D Systems Minneapolis, MN), CD64 (BioLegend San Diego, CA), CD68 (Invitrogen Waltham, MA), and CD163 (BD Pharmingen San Diego, CA) staining. Following initial characterization of basal and LPS induced cytokine and chemokine production, iPSC Line A was used for all subsequent experiments.

### Human intestinal organoid (HIO) generation

iPSC derived HIO were generated from the same CD donor A iPSC line used for macrophage generation and studies, as recently reported ^8^. Definitive hindgut endoderm was generated as reported.^8^ Induced cells were harvested and seeded on AggreWell^TM^400 Microwell Plate (StemCell Technologies Cambridge, MA) at 3.6 million cells in each well for 24 hours. On the following day, the derived spheroids were harvested and embedded in Matrigel (BD Biosciences, San Jose CA) and incubated in media containing recombinant 50ng/mL hEGF (R&D Systems Minneapolis, MN), 100ng/mL hNoggin (Shenandoah Warminster, PA), and 5% Respondin-2 (CCHMC-PSCF Cincinnati, OH) conditioned media. HIO were passaged and re-embedded in matrigel weekly, and then harvested for assays four weeks after embedding. We examined HIO under basal conditions and after 72-hour co-culture with 50,000 LPS (Sigma-Aldrich St. Louis, MO) primed (100 ng/mL) iPSC-derived macrophages per well. Select experiments also included exposure to 50 μM ETYA (Sigma-Aldrich St. Louis, MO) for 3-14 days. HIO single cell preps were generated as reported.^8^

### RNA Sequencing (RNASeq)

RNA was isolated from macrophages using the Qiagen AllPrep RNA/DNA Micro Kit (Qiagen, Germantown, MD). The macrophage global pattern of gene expression was determined using RNASeq on the Illumina platform.^8^ Reads were quantified by kallisto using Gencode v24 as the reference genome. Protein-coding genes with Transcripts per Million (TPM) above 1 in 20% of the samples were included in subsequent analyses. Fold change (FC) ≥1.5 and false discovery rate (FDR) < 0.05 was used to defined differentially expressed genes in R package DESeq2 version 1.24.0.^8^ Gene Set Enrichment Analyses (GSEA) was conducted using ToppGene and ToppCluster, and network visualization utilized Cytoscape.v3.0.217.

### Luminex assay

Macrophage production of selected cytokines and chemokines was measured using a custom Luminex assay or ELISA. TNFA, IL-1b, OSM, IL-10 and CCL7 were measured using a multiplex Luminex bead assay by R&D Systems (R&D Systems, Minneapolis, MN). CCL2 and TL1A were measured using human CCL2 (MCP-1) and human TL1A ELISA kits by R&D Systems (R&D Systems, Minneapolis, MN). SEMA3C was measured using the human SEMA3C ELISA kit by LS Bio (LS Bio, Shirley, MC).

### TaqMan Low Density Array (TLDA) Array

The AllPrep DNA/RNA Micro kit was used to prepare HIO RNA (Qiagen, Germantown, MD). SuperScript IV VILO Master Mix was used to prepare cDNA, and pre-amplification was performed as indicated using TaqMan PreAmp Master Mix (ThermoFisher, Waltham, MA). Custom TaqMan Low Density Array (TLDA) cards (ThermoFisher, Waltham, MA) were used to perform real-time PCR on a 7900HT Fast Real-Time PCR System (Applied Biosystems, Waltham, MA). Glyceraldehyde-3-phosphate dehydrogenase (GAPDH) expression was used for normalization of target gene expression. The Taqman Assay IDs for the PCR primers were as follows: *ACTA2* (Hs00426835_g1), *COL1A1* (Hs00164004_m1), *COL4A5* (Hs01012435_m1), *DUOX2* (Hs00204187_m1), *EPCAM* (Hs00901885_m1), *GAPDH* (Hs99999905_m1), *NOX4* (Hs01379108_m1), *TGFB1* (Hs00998133_m1), *TLR4* (Hs00152939_m1), and *VIM* (Hs00958112_g1).

### Histology

Hematoxylin and eosin (H&E) staining and sirius red immunohistochemistry (Picro Sirius Red Stain Kit, Abcam Cambridge, MA) of HIO sections was performed as reported.^8^ Sirius red stained slides were analyzed using polarized light microscopy in ImageJ. HIO collagen architecture was also assessed using whole mount staining for Vimentin (Invitrogen Carlsbad, CA), E-cadherin (R&D Systems Minneapolis, MN) and collagen1 (Abcam Waltham, MA). Immunofluorescence detection of CD14 and CD68 was performed using anti-CD14 antibody (Sigma-Aldrich St. Louis, MO) and CD68 antibody (Sigma-Aldrich St. Louis, MO), co-stained with DAPI (Thermo Fisher Scientific Waltham, MA). Immunofluorescence detection of alpha-smooth muscle actin (αSMA) and vimentin (VIM) co-stained with DAPI was performed as reported.^8^ An Olympus BX51 microscope with a DP80 camera or an Olympus BX43 with a DP74 camera were utilized for image acquisition. ImageJ (National Institutes of Health, Bethesda, MD) software was used to measure the surface area for these immune or stromal cell stains.

### Statistical Considerations

Data are shown as the mean (SEM). Data normality was tested using the Shapiro-Wilk test. The unpaired t test was used for normally distributed data, and the Mann-Whitney test for data which did not pass the normality test. A one-way ANOVA with Dunnett’s multiple comparisons tested differences between treatment groups for normally distributed data, while a Kruskal-Wallis test with Dunn’s multiple comparisons test was employed for data which did not pass the normality test in GraphPad PRISM 9.0.

### Ethical Statement

The patient-based studies were approved by the Institutional Review Board at Cincinnati Children’s Hospital Medical Center, and consent was obtained from parents and adult subjects and assent from pediatric subjects aged 11 and above.

## Results

### CD patient iPSC-derived macrophages exhibit a pro-inflammatory pattern of cytokine and chemokine production implicated in anti-TNF non-response and disease complications

Our collaborators previously reported an inflammatory macrophage:fibroblast gene expression module notable for up-regulation of *TNFA, IL1B, OSM,* and *CCL2* in the CD ileum.^1^ Patients with higher expression of this module at diagnosis were more likely to be anti-TNF non-responders and experience disease complications.^1,2,4^ Studies over the past decade have also implicated TL1A in both inflammatory and fibrotic pathways in CD, with anti-TL1A antibodies entering phase II trials.^12,13,14,15^ We therefore investigated if macrophages generated from iPSC lines from four pediatric-onset CD patients (iPSC lines A, B, C, D, Fig. 1) would produce these cytokines and chemokines upon LPS stimulation. LPS induced a significant increase in TNFA, IL1B, and OSM production by macrophages derived from each of the four CD iPSC lines.

**Figure 1.**
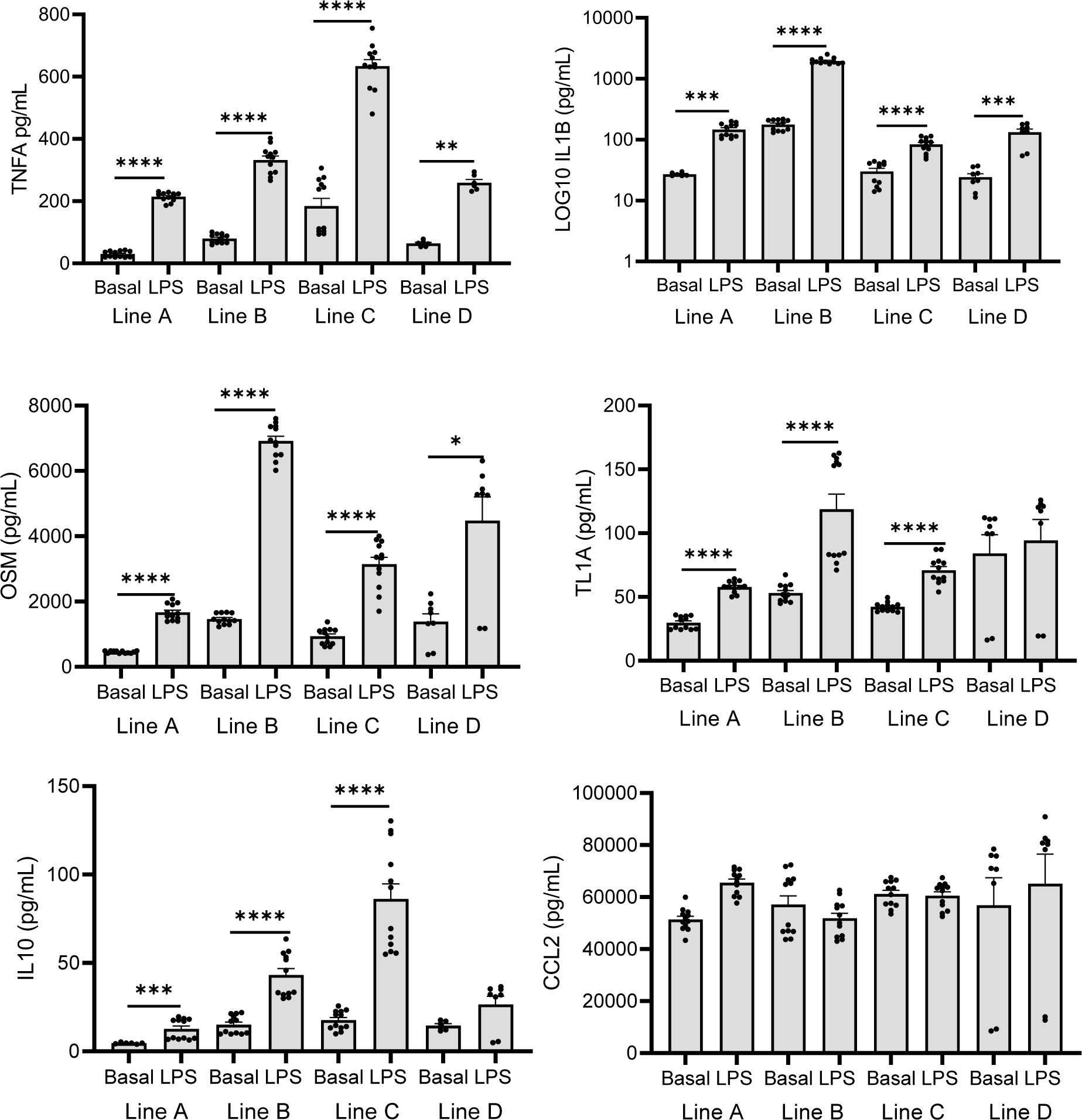
Cytokine and Chemokine Production by iPSC-derived Macrophages Stimulated with LPS. iPSC lines A, B, C, and D derived from PBMC obtained from four pediatric-onset CD patients were differentiated into macrophages. iPSC-derived macrophages were assayed under basal conditions or following LPS (100 ng/mL for 72 hours) stimulation. Cytokine and chemokine secretion was measured using a custom luminex assay or ELISA. Data are shown as the mean (SEM), n=6-12 per group, *p<0.05, ***p<0.001, ****p<0.0001.

TL1A and IL10 production was induced by LPS for each of the lines except for line D. The CCL2 chemokine was constitutively produced by macrophages derived from each of the four CD iPSC lines, without any further increase in response to LPS. These data confirmed that CD patient iPSC-derived macrophages produce clinically relevant cytokines and chemokines under basal and LPS stimulated conditions.

### CD patient iPSC-derived macrophages exhibit primary macrophage morphology and expression of pro-inflammatory cell surface proteins

As LPS stimulated cytokine production by macrophages derived from CD iPSC line A was comparable to the other three lines, macrophages and HIO derived from CD iPSC line A were used for all subsequent experiments. Cytospin confirmed typical primary macrophage morphology (Fig. 2A), with flow cytometry confirming that most of the cells expressed cell surface CD14 (Fig. 2B). While the frequency of CD14 expression did not vary with LPS or ETYA exposure, CD14 mean fluorescence intensity (MFI) increased with LPS and was suppressed with ETYA co-treatment (Fig. 2C). We observed a three-fold higher frequency of macrophages expressing CD68 relative to CD163; this did not vary across the groups (Fig. 2C). Consistent with a pro-inflammatory phenotype, 100% of the macrophages expressed CD64; this did not vary across groups (Fig. 2B and data not shown). However, we observed a significant reduction in CD64 cell surface abundance with ETYA exposure (Fig. 2C). These data demonstrated that iPSC-derived macrophages exhibited a pro-inflammatory pattern of cell surface markers, with suppression of the activation marker CD64 by ETYA

**Figure 2.**
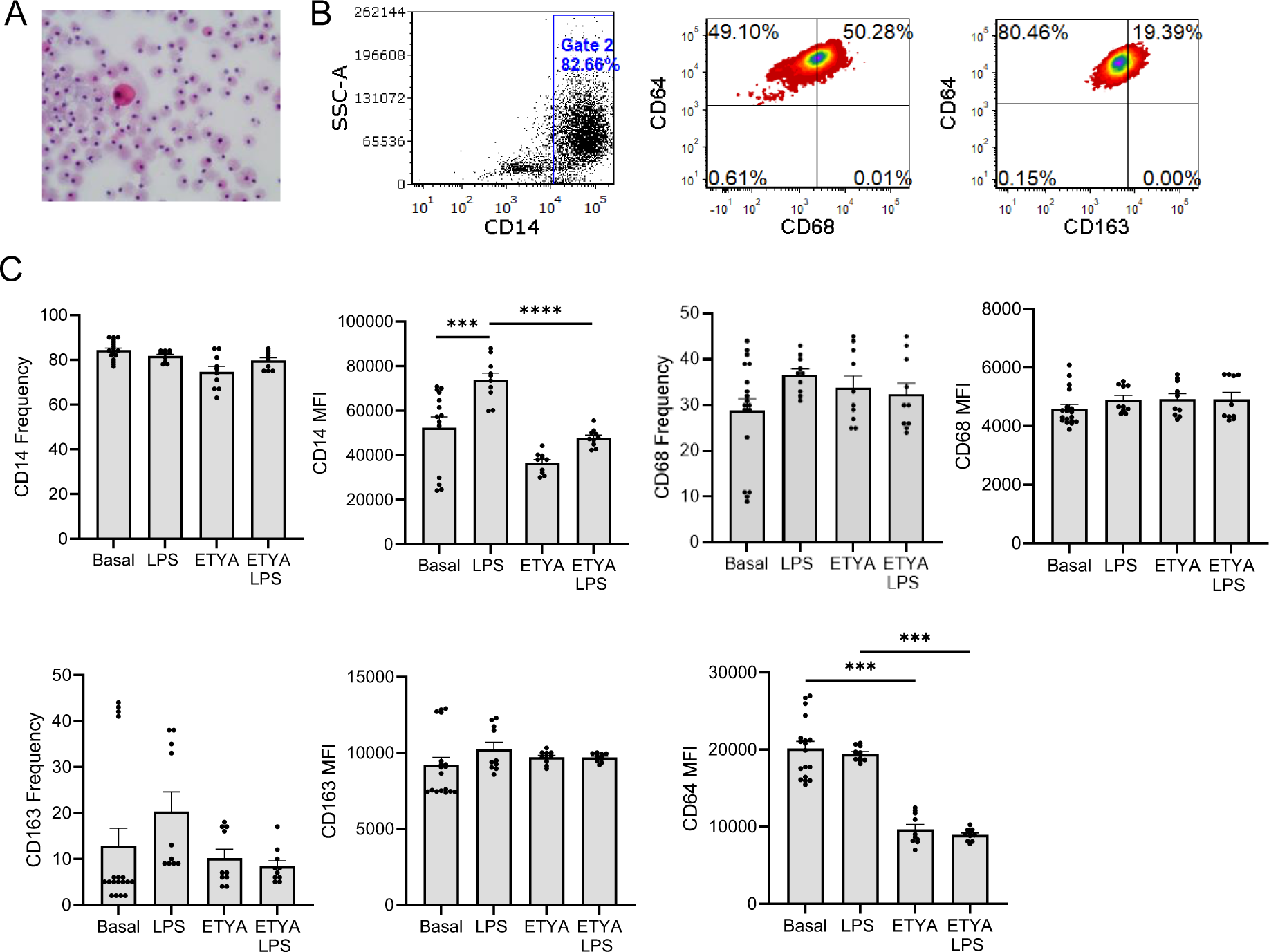
Morphology and Activation Marker Expression of iPSC-derived Macrophages Stimulated with LPS and ETYA. iPSC-derived macrophages were assayed under basal conditions or following LPS (100 ng/mL for 72 hours) ± ETYA (50 μM for 72 hours) stimulation. A) iPSC-derived macrophage morphology was assessed using cytospin. B) Representative scatter plots are shown for CD14, CD64, CD68, and CD163 staining. C) The frequency and MFI for CD14, CD64, CD68, and CD163 under the different conditions is shown. Data are shown as the mean (SEM), n=10-18 per group, ***p<0.001, ****p<0.0001.

### LPS primed CD patient iPSC-derived macrophages exhibit a pro-fibrotic global pattern of gene expression which is suppressed by ETYA

Since macrophages appeared to have an inflammatory signature, we therefore investigated if there was an increase in pro-fibrotic gene expression. Treatment with ETYA alone led to differential expression of 1062 genes, relative to basal conditions (Fig. 3A and Supplemental Table 1). As predicted by our recent study,^4^ this included up-regulation of genes regulating lipid metabolism (Reactome Metabolism of Lipids, FDR B&H: 1.08E-05), and target genes for the PPARG agonist rosiglitazone (FDR B&H: 7.42E-17). Stimulation with LPS alone led to differential expression of 1646 genes (Fig. 3A and Supplemental Table 2). Remarkably, LPS stimulated CD iPSC-derived macrophages were highly enriched for genes detected by scRNA-seq in inflammatory macrophages in the CD ileum^1^ (Fig. 3B & 3C, ToppCell Atlas, FDR B&H: 1.968E-121) and previously implicated via genetic analysis in host:microbe interactions involved in CD pathogenesis^12^ (FDR B&H: 2.19E-26). Most relevant to the current study, we also observed enrichment for up-regulated genes encoding the matrisome (FDR B&H: 2.16E-11). Co-treatment with ETYA and LPS resulted in differential expression of 859 genes, relative to stimulation with LPS alone (Fig.3A and Supplemental Table 3). This included suppression of genes expressed by small intestinal monocytes and myeloid cells (FDR B&H: 2.26E-45), including those encoding CD risk genes involved in host:microbe interactions (FDR B&H: 1.22E-11) and the matrisome^13^ (FDR B&H: 1.65E-08, Figs. 3B & 3C and Supplemental Table 3). These data showed that LPS-primed macrophages exhibited a global pattern of gene expression enriched in CD ileal inflammatory macrophages, with ETYA co-treatment inhibiting expression of pro-fibrotic genes linked to the matrisome.

**Figure 3.**
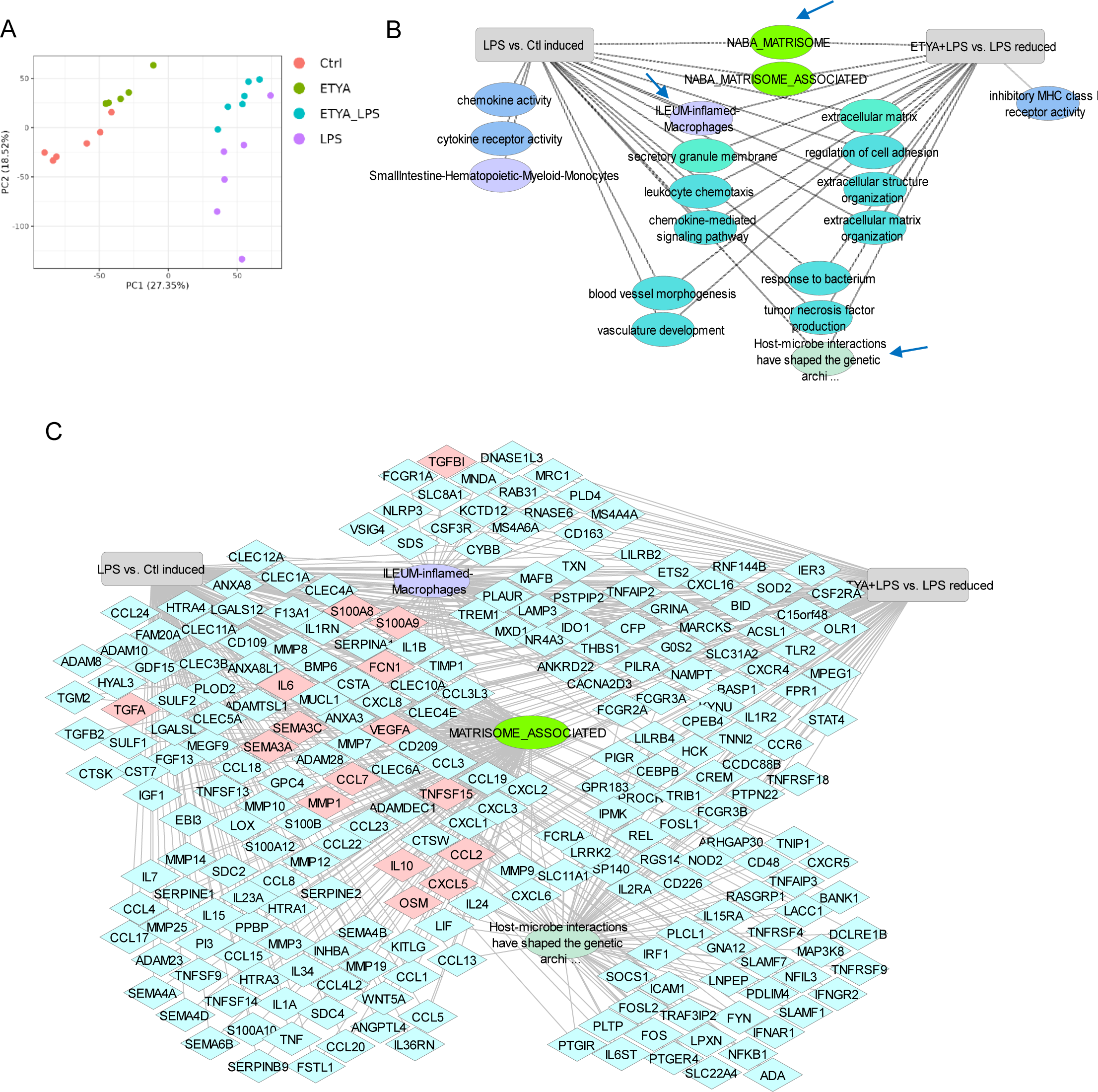
Differentially Expressed Genes and Pathways in iPSC-derived Macrophages Stimulated with LPS and ETYA. iPSC-derived macrophages were assayed under basal conditions or following LPS (100 ng/mL for 72 hours) ± ETYA (50 μM for 72 hours) stimulation. Differentially expressed genes were defined using bulk RNASeq (n=6 per group, with FDR<0.05 and fold change (FC) ≥1.5). A) Principal components analysis was used to visual variation in overall gene expression (11,141 genes that passed expression filtering) between the groups. ETYA_LPS: macrophages co-treated with ETYA and LPS for 72 hours. B) Functional annotation enrichment analysis of the 984 genes up regulated in LPS stimulated macrophages (LPS vs Ctl induced), and the 722 genes suppressed by ETYA co-treatment of LPS stimulated macrophages (ETYA+LPS vs LPS reduced), was completed using ToppGene, ToppCluster, and Cytoscape. Enriched signaling pathways, cell types, and biologic functions are as shown. Lines indicate shared or unique up- and down-regulated pathways between the two treatment conditions. C) Genes up-regulated by LPS stimulation, and down-regulated by ETYA co-treatment of LPS stimulated macrophages are shown relative to enrichment within ileal inflamed macrophages, the matrisome pathway, and/or host-microbe interactions implicated in CD genetic architecture. Genes specifically implicated in refractory CD and further tested in the current report are highlighted in pink.

### Eicosatetraynoic-acid (ETYA) regulation of LPS stimulated CD patient iPSC-derived macrophage cytokine and chemokine production

We therefore next asked whether these differences in gene expression would be reflected in differences in cytokine or chemokine production, with a focus upon mediations previously implicated in ant-TNF response and disease complications.^1,2,4,14,15^ More recently, *SEMA3C* which we found was differentially expressed in the gene expression data set has also been implicated in wound healing in the gut^16^, and development of liver fibrosis.^17^ We therefore tested for variation in macrophage production of these cytokines and chemokines with LPS and ETYA stimulation. LPS induced macrophage TNFA, IL1B, OSM, TL1A, CCL7, CXCL5, and SEMA3C secretion, with relatively lower levels of IL10 production (Fig. 4). We did not observe macrophage production of the LAP complex containing TGFB1 under any of the conditions tested (data now shown). CCL2 was constitutively produced at a high level. ETYA co-treatment did not reduce inflammatory cytokine production, but did inhibit production of the CCL2, CCL7, and CXCL5 chemokines implicated in monocyte and neutrophil recruitment to the gut.^1^ Interestingly, SEMA3C was induced by ETYA alone; this did not vary further with ETYA/LPS co-treatment. These data demonstrated that iPSC-derived macrophages exhibit a pro-inflammatory pattern of cytokine and chemokine production implicated in anti-TNF non-response, which was partially inhibited specifically for chemokines by ETYA.^1,4^

**Figure 4.**
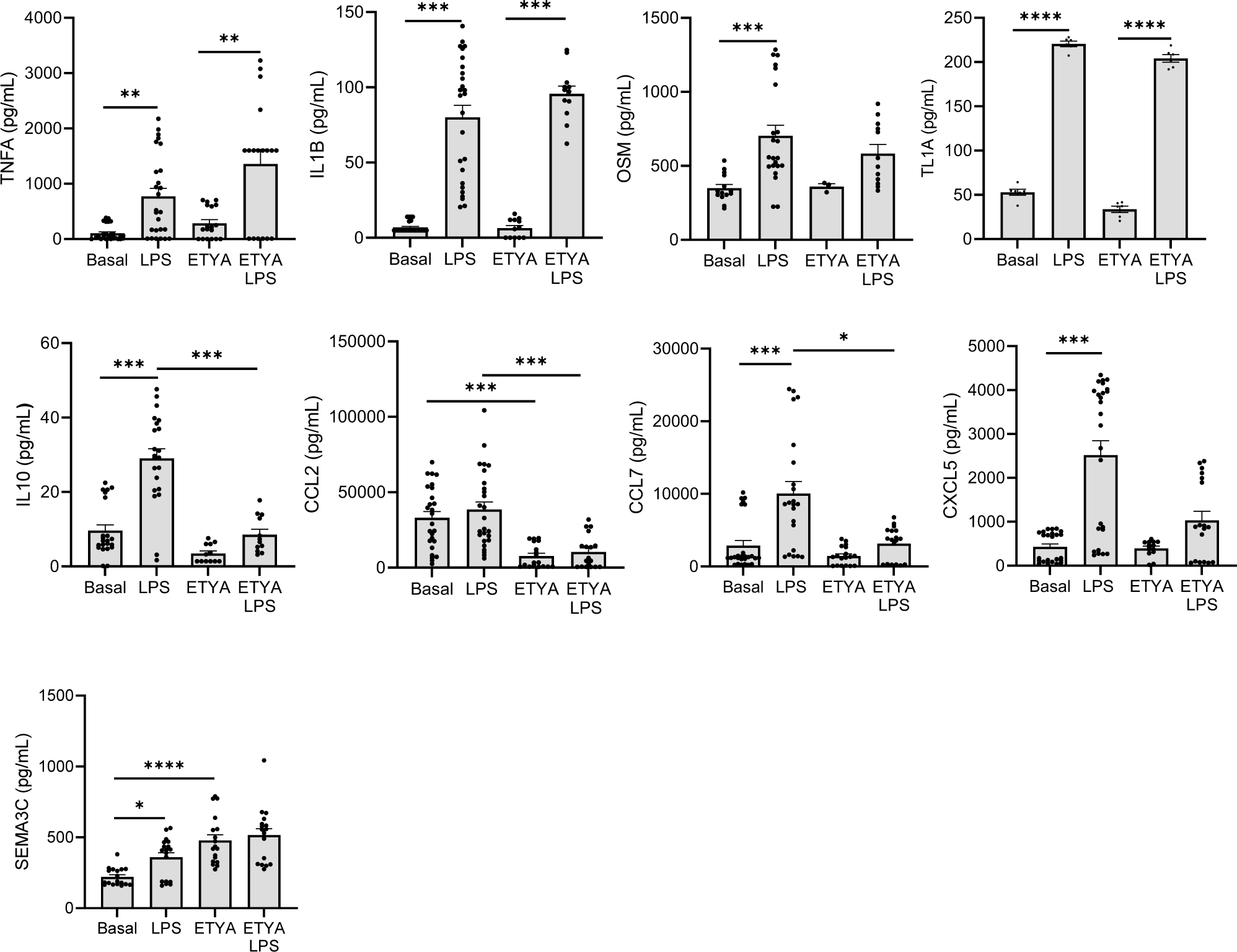
Cytokine and Chemokine Production by iPSC-derived Macrophages Stimulated with LPS and ETYA. iPSC-derived macrophages were assayed under basal conditions or following LPS (100 ng/mL for 72 hours) ± ETYA (50 μM for 72 hours) stimulation. Cytokine and chemokine secretion was measured using a custom luminex assay or ELISA. Data are shown as the mean (SEM), n=18-21 per group, *p<0.05, **p<0.01, ***p<0.001, ****p<0.0001.

### ETYA reduces the frequency of LPS stimulated CD14+ myeloid cells and ACAT2+ stromal cells in the iPSC-derived macrophage:HIO co-culture system

We therefore next tested effects of ETYA in the setting of co-culture of LPS primed macrophages with HIO. HIO growth and morphology did not vary with LPS primed macrophage co-culture (Mac), treatment with ETYA alone, or ETYA treatment prior to LPS primed macrophage co-culture (ETYA/Mac, Fig. 5A and data not shown). As expected, we observed an increase in the frequency of CD14+ and C68+ myeloid cells following co-culture of LPS primed macrophages with HIO (Figs. 5B & 5D). The frequency of CD14+ cells was reduced by ETYA pre-treatment, consistent with effects of ETYA upon isolated macrophages. Consistent with the pro-fibrotic effects of LPS stimulation upon isolated macrophages, co-culture of LPS primed macrophages with HIO increased the frequency of cells expressing myofibroblast (ACTA2) and fibroblast (VIM) cell surface markers (Figs. 5C & 5D). The frequency of both ACAT2+ and VIM+ stromal cells was reduced by ETYA pre-treatment. These data demonstrated anti-inflammatory and anti-fibrotic effects of ETYA pre-treatment upon key myeloid and stromal cell populations in the LPS primed macrophage:HIO co-culture system.

**Figure 5.**
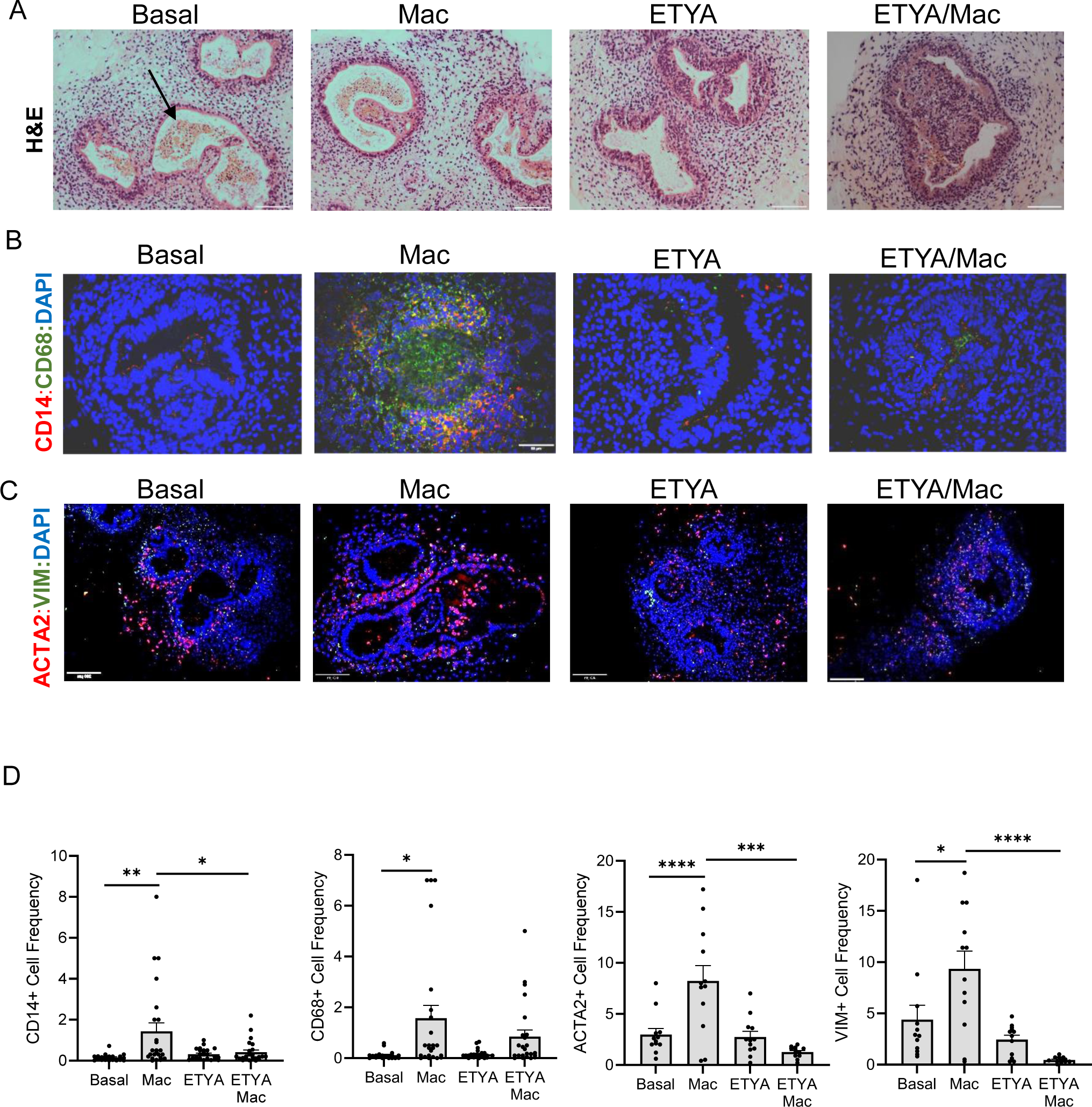
Human Intestinal Organoid Morphology and Macrophage and Fibroblast Abundance. Human intestinal organoids were assayed under basal conditions, following co-culture with LPS stimulated (100 ng/mL) macrophages for 72 hours (Mac), following ETYA pre-treatment (50 uM) for 12 days (ETYA), and following ETYA pre-treatment and LPS stimulated macrophage co-culture (ETYA/Mac). A) HIO morphology was assessed using H&E staining. The HIO lumen is indicated by the arrow. B) The frequency of CD14 positive (red) and/or CD68 positive (green) monocytes and macrophages is shown. Nuclei were stained with DAPI (blue). C) The frequency of human intestinal organoid (HIO) alpha smooth muscle actin (ACTA2) positive (red) and/or vimentin (VIM) positive (green) stromal cells is shown. Nuclei were stained with DAPI (blue). Representative images are shown. D) The frequency of CD14+, CD68+, ACAT2+, and VIM+ cells was determined using ImageJ. Data are shown as the mean (SEM), n=20-24 per group, *p<0.05, **p<0.01, ***p<0.001, ****p<0.0001.

### Eicosatetraynoic-acid (ETYA) prevents pro-fibrotic effects of iPSC-derived macrophages upon HIO gene expression and collagen content

Finally, we tested whether co-culture of LPS primed macrophages with HIO would induce pro-fibrotic gene expression, and whether this would be regulated by ETYA. Neither expression of the epithelial marker EPCAM, nor the LPS receptor TLR4, varied significantly across the groups (Fig. 6). Interestingly the epithelial NADPH oxidase DUOX2 required for ileal homeostasis was induced by ETYA.^18-21^ Expression of the pro-fibrotic *TGFB1* gene was induced with LPS primed macrophage co-culture; this was not inhibited by ETYA pre-treatment (Fig. 6). Macrophage co-culture also induced HIO expression of alpha smooth muscle actin (*ACTA2*) and *NOX4* expressed by activated myofibroblasts, and *COL1A1* encoding type I collagen (Fig. 6). These pro-fibrotic changes in gene expression were prevented by ETYA pre-treatment, together with suppression of vimentin (VIM) expressed by fibroblasts. Under these conditions the expression of *COL4A5* did not vary. These data confirmed induction of pro-fibrotic genes expressed by HIO under LPS-primed macrophage co-culture conditions, which was largely inhibited by ETYA pre-treatment. The sup-epithelial collagen band detected in HIO is shown in Figs. 6B and 6C. LPS primed iPSC-derived macrophages and HIO were co-cultured for three days. An increase in HIO type I and type III collagen content was observed following LPS primed macrophage co-culture (Fig. 6D). HIO collagen content under macrophage co-culture conditions was reduced by ETYA pre-treatment for 12 days (Fig. 6D). These data demonstrated that ETYA reduced HIO extra-cellular matrix gene expression and collagen content under inflammatory macrophage co-culture conditions, consistent with our recent report in HIO alone.^8^

**Figure 6.**
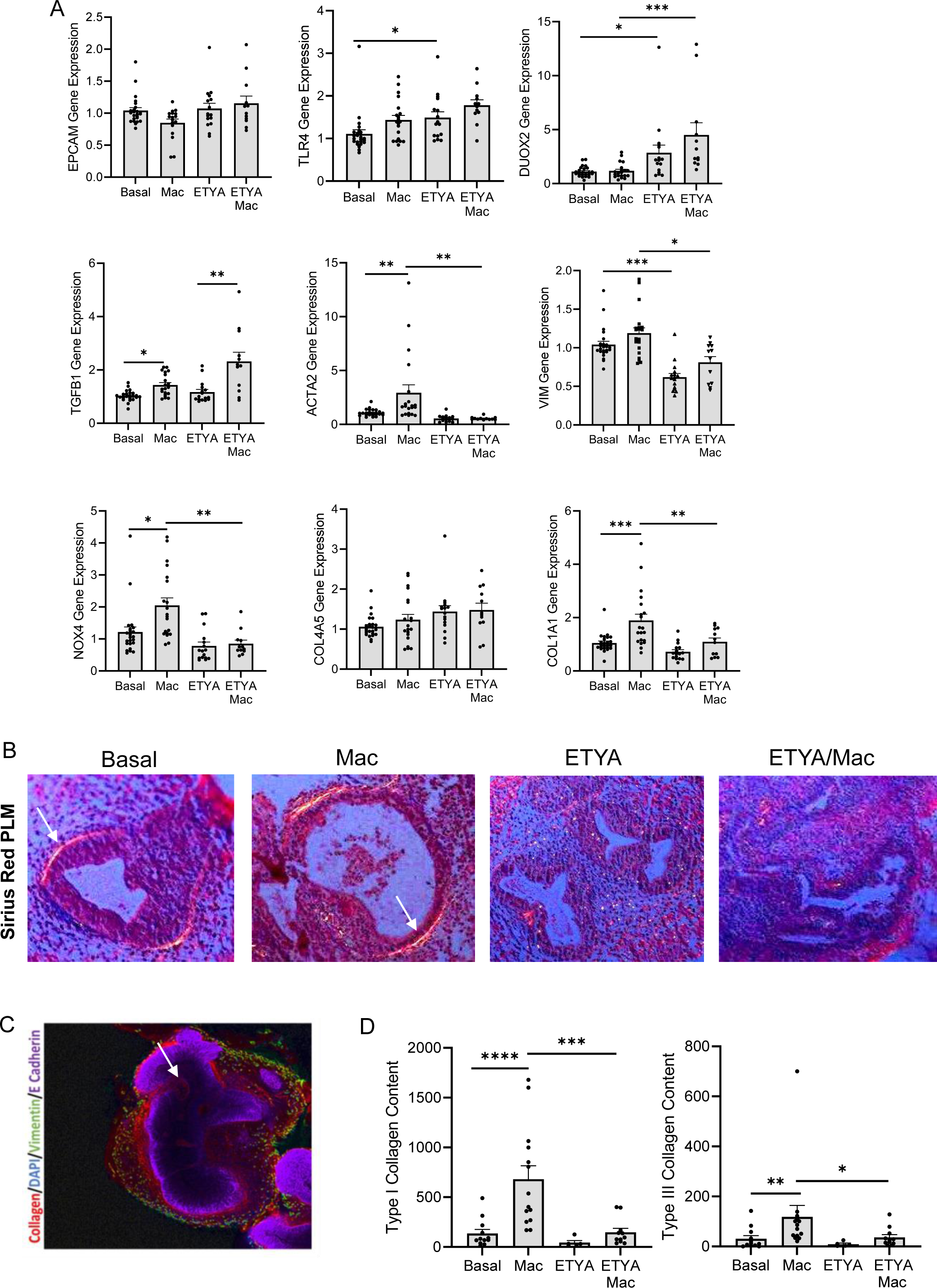
Variation in Human Intestinal Organoid Gene Expression and Collagen Content with Macrophage Co-Culture and ETYA Pre-treatment. Human intestinal organoids (HIO) were assayed under basal conditions, following co-culture with LPS stimulated (100 ng/mL) macrophages for 72 hours (Mac), following ETYA pre-treatment (50 uM) for 12 days (ETYA), and following ETYA pre-treatment and LPS stimulated macrophage co-culture (ETYA/Mac). A) HIO gene expression was determined using a custom TLDA array. Data are shown as the mean (SEM), n=12-24 per group, *p<0.05, **p<0.01, ***p<0.001. B) HIO collagen content was determined using sirius red staining with polarized light microscopy (PLM). Representative images are shown, with the sub-epithelial collagen band shown by the arrow. C) Whole mount staining was used to define the sub-epithelial collagen band (red) relative to the organoid structure (E Cadherin, purple, with lumen indicated by the arrow), and surrounding vimentin positive fibroblasts (green). D) Type I and Type III collagen content was measured using sirius red staining with polarized light microscopy (PLM) in ImageJ. Data are shown as the mean (SEM), n=5-14 per group, *p<0.05, **p<0.01, ***p<0.001.

## Discussion

Biologics targeting anti-TNF are the mainstay of pediatric Crohn’s Disease (CD) therapy.^22^ However, anti-TNF non-response, and progression to fibrotic end-organ injury still complicates the course of many patients.^23-25^ Our perturbagen analysis of the treatment naïve pediatric CD ileal transcriptome utilized a dataset containing over 2000 small molecules to identify compounds likely to reverse the ileal gene signature for anti-TNF non-response and subsequent stricturing.^4^ These data prioritized studies of these candidate small molecules in a relevant patient-derived model system, to advance personalized medicine and discovery of new therapeutic approaches. A critical barrier to progress has been the lack of a patient-derived model system to test candidate small molecules predicted to regulate mechanisms driving anti-TNF response. Our proposed studies have begun to address this barrier, by testing the novel concept that iPSC lines prepared from CD patients will yield a macrophage:organoid co-culture system which will capture critical inflammatory responses linked to anti-TNF response.^5-7,10,26^ We found that LPS stimulated iPSC-derived macrophages exhibited a global pattern of gene expression and cytokine production highly relevant to refractory CD.^1-4^ A small molecule prioritized by our perturbagen analysis, ETYA, exerted key anti-inflammatory and anti-fibrotic effects.^4,8^

CD patient iPSC-derived human intestinal organoids (HIO) co-cultured with iPSC-derived macrophages are an attractive experimental model for human CD. ^5-7,26^ HIO derived from iPSC contain epithelial cells and fibroblasts and increase collagen production in response to TGFβ.^5-7^ We found that LPS stimulated iPSC-derived macrophages expressed a global pattern of gene expression enriched in CD ileal inflammatory macrophages,^1^ and induced a pro-fibrotic gene program with collagen production in HIO co-culture. Retention of somatic memory in iPSC lines derived from CD PBMC would be anticipated to minimal, with donor genetic architecture controlling epigenetic and transcriptional variation in subsequent differentiated cell types.^27^ We anticipate that iPSC-derived macrophages and HIO will therefore capture the integrated genetic output driving variation in cell function linked to inflammatory responses and tissue collagen content.^28^ Future studies will characterize macrophages and HIO derived from a wider range of CD iPSC lines, stratified by both genetic architecture and clinical phenotype.^1,7,8,12^

The synthetic long chain fatty acid eicosatetraynoic acid (ETYA) was prioritized by our perturbagen analysis, and predicted to both inhibit macrophage and fibroblast activation, in part via PPAR signaing.^4^ Consistent with this, ETYA induced PPAR dependent lipid metabolic pathways and suppressed macrophage CD64 expression which we have previously linked to increased anti-TNF drug clearance and CD endoscopic activity.^29,30^ While ETYA inhibited macrophage production of several pro-inflammatory chemokines which drive gut recruitment of monocytes and neutrophils, it did not inhibit production of key cytokines including TNFA, IL1B, OSM, or TL1A implicated in fibrostenosis.^1,14,28^ Nevertheless, ETYA pre-treatment of HIO inhibited pro-fibrotic effects of co-cultured macrophages. This may be due to inhibition of alternate pro-fibrotic macrophage products; it will be important to test this more broadly in a future study. Alternately, based upon prior studies, ETYA pre-treatment likely inhibited fibroblast activation and collagen production in response to a variety of macrophage cytokines.^4^ Future studies of isolated patient-specific fibroblasts will be required to test this.

Our study has several strengths, but also some limitations. We found that iPSC-derived macrophages provide a relevant model system to test mechanisms driving CD ileal inflammatory macrophages, and that HIO collagen content can be modulated with both macrophage signals and candidate small molecules.^31^ However, in the tissue culture system, this consists primarily of a sub-epithelial collagen band.^5-8^ Our collaborators have developed transplantation of the HIO under the kidney capsule of immune deficient mice, to address this limitation.^32^ Under these conditions, over 12 weeks the full sub-mucosa and smooth muscle layers of the intestine develop, recapitulating the collagen architecture of CD strictures.^26,32,33^ It recently been reported that co-cultured macrophages may be stably incorporated into the transplanted HIO under these conditions.^26^ It will be important in future studies to take this approach, to test whether specific CD patient genetic profiles, and candidate small molecules including ETYA, exert significant effects upon tissue architecture in the murine iPSC-derived macrophage:HIO transplantation system.

Derivation of iPSC from PBMC eliminates confounding epigenetic effects within primary tissues, providing an integrated model of genetic effects linked to important clinical phenotypes including anti-TNF response. While the current study provides an important proof of concept for this approach, in the future a wider range of iPSC lines from males and females, and patients with diverse racial and ethnic backgrounds, will need to be tested to determine whether macrophage:HIO endpoints vary with sex or race. Ideally this will include CRISPR/Cas9 gene editing of key candidate genes to provide isogenic controls across diverse patient backgrounds.^34^ Along these lines a member of our group has recently developed a novel model of metabolic dysfunction-associated steatohepatitis (MASH) with inflammatory and fibrotic features highly relevant to the current study.^35^ This included the first population-level survey of MASH pathogenesis in human liver organoids (HLO) derived from 24 patients with a diverse racial and ethnic background. The current data set supports a follow-up CD study of similar scale, with application of HIO as recently reported to mechanisms of epithelial injury and fibrosis relevant to Crohn’s Disease.^36^

In conclusion, we found that CD patient iPSC-derived macrophages provide a highly relevant model system for CD ileal inflammatory macrophages, with production of key inflammatory and fibrotic mediators implicated in a refractory disease.^1-4^ LPS primed macrophages exerted pro-fibrotic effects when co-cultured with isogenic HIO. As proof of concept for the perturbagen bioinformatic approach, a candidate small molecule exerted anti-inflammatory effects upon LPS primed macrophages and prevented HIO pro-fibrotic responses to co-cultured macrophages.^4^ These data and model system will advance our ability to perform screens of candidate therapies, advancing personalized medicine in CD.

## Acknowledgements

Mary Kate Ewalt, Amanda Schreibeis. Matthew Kofron, and Christopher Mayhew provided outstanding technical support. Luminex assays and CCL2 ELISA were performed by the Research Flow Cytometry Facility in the Division of Rheumatology at Cincinnati Children’s Hospital Medical Center.

## Abbreviations

ACTA2: smooth muscle actin alpha 2
B1: inflammatory behavior
B2: stricturing behavior
B3: internal penetrating behavior
CCL2: C-C Motif Chemokine Ligand 2
CCL7: C-C Motif Chemokine Ligand 7
CD: Crohn’s Disease
COL1A1: Collagen Type I Alpha 1 Chain
COL4A5: Collagen Type IV Alpha 5 Chain
Ctrl: control
DAPI: 4′,6-diamidino-2-phenylindole
CXCL5: C-X-C Motif Chemokine Ligand 5
DUOX2: Dual Oxidase 2
ECM: extra-cellular matrix
ECM1: Extracellular Matrix Protein 1
EGF: epidermal growth factor
EPCAM: epithelial cell adhesion molecule
ETYA: eicosatetraynoic acid
H&E: hematoxylin and eosin stain
HIO: Human Intestinal Organoid
IBD: Inflammatory Bowel Disease
IL: interleukin
INFL: inflammatory
iPSC: induced pluripotent stem cell
IQR: inter-quartile range
LAP: latency-associated peptide
LPS: lipopolysaccharide
MMP: mitochondrial membrane potential
L1: ileum-only location
L2: colon-only location
L3: ileo-colonic location
Mac: macrophage
MFI: mean fluorescence intensity
NADPH: nicotinamide-adenine dinucleotide phosphate
NOX4: NADPH oxidase 4
PBMC: peripheral blood mononuclear cell
PCA: principle components analysis
PLM: polarized light microscopy
PPAR: peroxisome proliferator-activated receptor
OSM: oncostatin M
QC: quality control
REF: reference
RESP: respiratory
RNASeq: RNA sequencing
ROS: reactive oxygen species
SEMA3C: semaphorin 3C
SSC-A: side scatter
TGFB: transforming growth factor beta
TGS: targeted gene sequencing
TL1A: TNF Ligand-Related Molecule 1
TLDA: TaqMan Low Density Array
TNF: tumor necrosis factor
VIM: vimentin

## Data deposition

The human iPSC-derived macrophage RNASeq data are deposited in GEO under GSE248009

## Writing assistance

Not applicable

## Author Contributions

Study concept and design: TT, YH, JW, LAD

Acquisition of data: IJ, BD, EA, EB, KI

Analysis and interpretation of data: IJ, BD, EA, EB, TB, YH, LAD

Drafting of the manuscript: IJ, BD, LAD

Critical revision of the manuscript for important intellectual content: KT, TT, JW, LAD

Obtained funding: BD, TT, YH, JW, LAD

Administrative, technical, or material support: IJ, EA, EB, KT, KI

Study supervision: IJ, LAD

## Conferences

Presented in part at the 2023 Digestive Diseases Week annual meeting, Chicago, IL, USA.

